# Deep Local Analysis evaluates protein docking conformations with Locally oriented Cubes

**DOI:** 10.1101/2022.04.05.487134

**Authors:** Yasser Mohseni Behbahani, Simon Crouzet, Elodie Laine, Alessandra Carbone

## Abstract

With the recent advances in protein 3D structure prediction, protein interactions are becoming more central than ever before. Here, we address the problem of determining how proteins interact with one another. More specifically, we investigate the possibility of discriminating near-native protein complex conformations from incorrect ones by exploiting local environments around interfacial residues. Deep Local Analysis (DLA)-Ranker is a deep learning framework applying 3D convolutions to a set of locally oriented cubes representing the protein interface. It explicitly considers the local geometry of the interfacial residues along with their neighboring atoms and the regions of the interface with different solvent accessibility. We assessed its performance on three docking benchmarks made of half a million acceptable and incorrect conformations. We show that DLA-Ranker successfully identifies near-native conformations from ensembles generated by molecular docking. It surpasses or competes with other deep learning-based scoring functions. We also showcase its usefulness to discover alternative interfaces.

**Availability:** http://gitlab.lcqb.upmc.fr/dla-ranker/DLA-Ranker.git

## 1 Introduction

Protein-protein interactions play a central role in virtually all biological processes. Reliably predicting who interacts with whom in the cell and in what manner would have tremendous implications for bioengineering and medicine. Hence, a lot of effort has been put into the development of methods for simulating protein-protein docking [26, 27]. While highly efficient algorithms can exhaustively sample the space of complex candidate conformations [40], correctly evaluating and ranking these conformations remains challenging.

The classical docking and scoring paradigm has been recently challenged by the spectacular advances in protein structure prediction with AlphaFold version 2 (AF2) [20] and RosettaFold [2]. In particular, a handful of studies have showcased the potential of AF2, or a slightly modified version, in fold-and-dock strategies [19, 14, 33,4]. Nevertheless, they have also emphasized clear limitations. AF2 performs poorly on some eukaryotic complexes, antibody-antigen complexes, and complexes displaying small interfaces [14, 4]. In such cases, the output is limited to an unreliable conformation. By contrast, docking algorithms allow for the generation of conformational ensembles useful to guide the prediction of interfaces, to gain insight into protein sociability [25], and to discover alternative binding modes and new partners [9]. These observations motivate the development of accurate and efficient methods assessing the quality of docking conformations.

The Critical Assessment of PRedicted Interactions (CAPRI) classifies predicted protein complex conformations in four categories, namely incorrect, acceptable, medium, and high-quality, based on the extent to which they differ from the corresponding experimental structures [28]. Recently, several methods leveraging deep learning have been proposed to discriminate near-native (acceptable or higher quality) from incorrect conformations [38, 49, 48, 5, 13]. They adopt a “global” perspective by assessing the quality of the full interface [38, 49, 48] or even the complex as a whole [13]. Standard 3D-convolutional neural networks (3D-CNN) have been applied to a voxelized 3D grid representing the entire interface [38, 49]. This representation has two limitations. First, when a fixed-size cube is used as grid, it might not cover very large and/or discontinuous interfaces. Using a very large cube to accommodate any interface is memory inefficient. Large cubes of fixed-size may also hinder the accuracy in case of small interfaces due to the information vanishing after a few layers of pooling. Second, since the 3D-CNN does not benefit from the rotational symmetry endowed to the Euclidean space, it is sensitive to the orientation of the candidate conformation and its output may change upon rotation of the input in an uncontrolled fashion.

Rotational data augmentation was used in [38] to limit this effect, but at the expense of dramatically increasing the computational cost for training the model. A more efficient solution is to use a SE(3)-equivariant CNN architecture instead of standard CNN. SE(3)-equivariant CNN make use of spherical harmonics, a set of functions defined on the unit sphere, to guarantee that a rotation of the input results in the same rotation of the output [6, 50, 16, 44]. In [13], SE(3)-equivariant hierarchical convolutions were applied to a point-cloud representation of the whole conformation. Finally, graph-based representations, as those used in GNN-DOVE [48] and DeepRank-GNN [41], are invariant to 3D rotations, but at the expense of losing information about the orientations of the atoms with respect to each other.

Alternatively, one can leverage the specific properties of proteins, whose building blocks (the amino acid residues) share the same chemical scaffold, to derive a SE(3)-equivariant representation. In single protein structure prediction, Ornate [36], Sato-3DCNN [43], and more recently AlphaFold2 [20], benefit from these properties and make use of oriented local frames centered on each protein residue. Such representation circumvents the problem of 3D rotational symmetry without the need for rotational data augmentation nor for SE(3)-equivariant convolutional filters.

In this work, we investigate the possibility of discriminating near-native complex conformations from incorrect ones by exploiting and combining two kinds of information: (i) local 3D-geometrical and physico-chemical environments around interfacial residues and (ii) regions of the interface with different solvent accessibility. We represent the interface by the unique and well-determined set of locally oriented residue-centred cubes lying between the interacting proteins (**Fig. 1A**). The cubes are oriented by defining local frames based on the common chemical scaffold of amino acid residues in proteins. A cube encapsulates the local environment of the residue, i.e. the local geometry of the residue together with its neighboring atoms. No evolutionary information associated to residues is considered. Our motivations for such a representation are multiple:

1. The number of known protein-protein complex structures is fairly limited. Breaking down these structures into interfacial residue-centred local environments allows training on a much larger set of input samples (cubes) compared to the number of interfaces.
2. Our representation guarantees that the output is invariant to the global orientation of the input conformation while fully accounting for the relative orientation of a residue with respect to its neighbours.
3. We wanted to investigate the minimal unit of information at the interface which is necessary to predict the quality of an interaction. By relying on minimal units, *i.e*. residue-centred cubes, one can also evaluate interfaces between three or more proteins.
4. The set of cubes belonging to the interface can be organized in three subsets depending on the solvent accessibility of the interfacial residues. The cubes within each subset are independent from each other and from the geometry of the surface. We wanted to study the contribution of these three subsets in ranking docking conformations.

**Figure 1:**
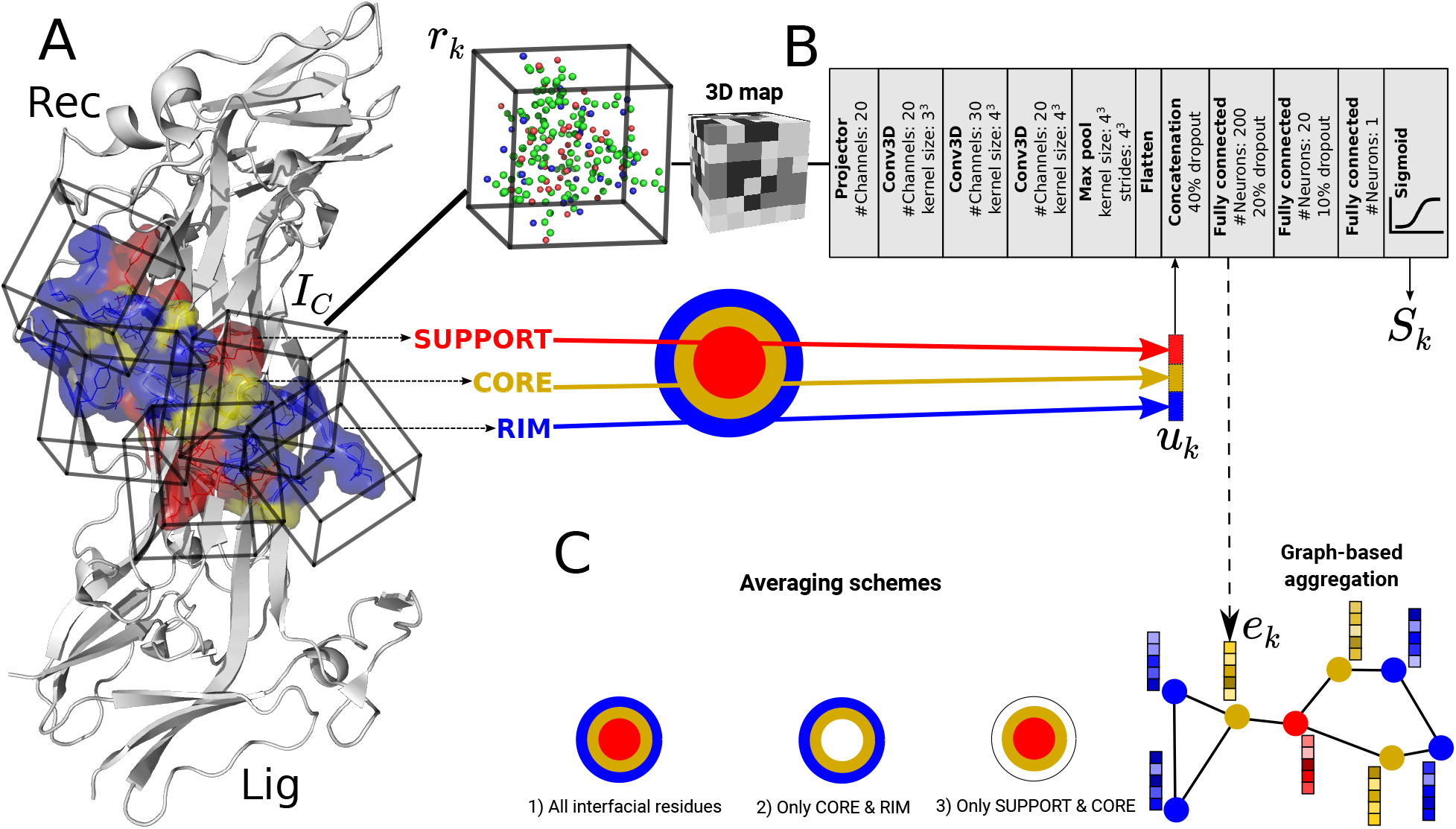
Interface representation and DLA-Ranker architecture. **A.** Representation of a protein interface as an ensemble of cubes (*I_C_*). Each cube (*r_k_* ∈ *I_C_*) is centered and oriented on an interfacial residue. It contains atoms belonging to the residue and its local environment (Carbon: green, Oxygen: red, Nitrogen: blue, Sulfur: yellow). A cube is labelled as being part of the Support (red), Core (gold), or Rim (blue) of the interface (one-hot encoded vector *u_k_*). **B.** Architecture of DLA-Ranker neural network. For input cube *r_k_* The network has two outputs: Score *S_k_* and embedding vector *e_k_*. **C.** The evaluation of the interface either by global averaging the local scores *S_k_* (1) over all interfacial residues, (2) over residues from SC, (3) over residues from CR, or by extracting embedding vectors *e_k_* and combining them through graph-based aggregation.

We propose Deep Local Analysis (DLA)-Ranker, a deep learning-based approach ranking candidate complex conformations by applying 3D-CNN to a set of locally oriented cubes representing the residues of the protein complex interface.

## 2 Methods

Our goal is to design a classifier that can effectively distinguish near-native protein candidate conformations from incorrect ones by learning from a local representation of the structure of the interface. Such representation should account for the local geometrical arrangement of interfacial atoms in the Euclidean space and their physico-chemical properties.

### 2.1 Protein-protein interface representation

DLA-Ranker takes as input a cubic volumetric map centred and oriented on each interfacial residue (**Fig. 1A**). Any residue displaying a change in solvent accessibility upon complex formation is considered as part of the interface. We used NACCESS [18] with a probe radius of 1.4 Å to compute residue solvent accessibility. To build the map, we adapted the method proposed in [36]. The atomic coordinates of the input conformation are first transformed to a density function. The density *d* at a point 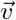 is computed as

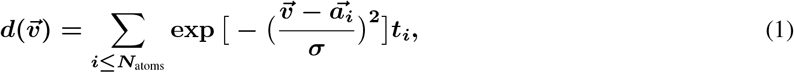

where 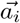 is the position of the *ith* atom, *σ* is the width of the Gaussian kernel and is set to 1Å, and *t_i_* is a vector of dimension 169 encoding some characteristics of the protein atoms. Namely, the first 167 dimensions correspond to the atom types that can be found in amino acids (without the hydrogens) [36], and the 2 other dimensions correspond to the two partners, the receptor and the ligand. Then, the density is projected on a 3D grid comprising 24 × 24 × 24 voxels of side 0.8Å. For the *nth* residue, the 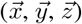 directions and the origin of the map are defined by the position of the atom N_*n*_, and the directions of C_*n*–1_ and *C_αn_* with respect to *N_n_* [36]. Thanks to this local frame definition, the map not only is invariant to the candidate conformation initial orientation but also provides information about the atoms and residues relative orientations.

Depending on the location of the residues at the interface, their geometrical and physico-chemical environments are expected to be very different. For instance, the map computed for a residue deeply buried in the interface will be much more dense than that computed for a partially solvent-exposed residue at the rim. This motivated us to explicitly give interfacial residues in three structure classes, the Support (S), the Core (C), and the Rim (R) (**Fig. 1A**), as defined in [29]. We one-hot encode the input residue class in a vector *u* and append it to the embedding computed by DLA-Ranker (see below and **Fig. 1B**-concatenation layer). The SCR classification previously proved useful for the prediction and analysis of protein-protein and protein-DNA interfaces [24, 37, 7].

### 2.2 DLA-Ranker architecture

The DLA-Ranker architecture comprises a projector, three 3D convolutional layers, a max pooling layer, and three fully-connected layers (**Fig. 1B**). The projector maps the feature vector of each voxel into a vector of size 20. Each convolutional layer is followed by a batch normalization layer. The max pooling layer exploits scale separability by preserving essential information of the input during coarsening of the underlying grid. The one-hot encoded vector of the residue structure class (*u*) is concatenated to the embedding derived from the convolutional layers (*i.e*. output of the flatten layer). To avoid overfitting, we used 40%, 20%, and 10% dropout regularization on the input, first and second layers of the fully-connected subnetwork, respectively. The last activation function (Sigmoid) outputs a score comprised between 0 and 1 for each input interfacial residue. The loss function is the binary cross-entropy measuring the difference between the probability distribution of the predicted output and the given label (0 or 1). The objective of training is to minimize this loss with respect to the trainable parameters: reaching higher output scores for the residues belonging to a near-native conformation and lower output scores for the residues of incorrect conformations. We used the Adam optimiser with a learning rate of 0.001 in TensorFlow [1].

### 2.3 Aggregation of individual residue-based scores

To evaluate a candidate conformation, DLA-Ranker applies global averaging on the individual residue scores over the interface. The predicted quality *Q* of conformation *C* is expressed as

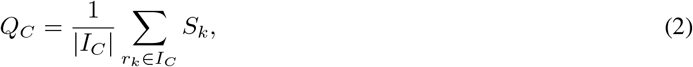

where *I_C_* is the ensemble of interfacial residues and *S_k_* is the score predicted by the network for the input 3D map centred on residue *r_k_*.

To investigate whether we could improve on this global averaging baseline, we considered two approaches. First, we proposed two additional evaluation schemes based on an average restricted to a selection of subsets of residues at the interface: (i) residues of S and C regions and (ii) residues of C and R regions (**Fig. 1C**). Second, we applied different weights to the residues comprising the interface by using graph-based attention [46] (**Fig. 1C** and **Fig. SI 1**). Namely, we extracted the embeddings *e_k_* computed by the first fully connected layer of DLA-Ranker and used them as node features in a graph representing the interface, where two nodes are linked if the distance between their associated residues is less than 5.0 Å. We apply one layer of self-attention and predict a unique score estimating the quality of the whole interface (**Fig. SI 1**).

### 2.4 Datasets

To train and test DLA-Ranker and compare its performance with different approaches we used three databases of docking conformations.

#### 2.4.1 CCD4PPI: docking conformations produced by MAXDo

We compiled our primary database, which we call CCD4PPI, from two complete cross-docking experiments performed on the datasets P-262 [11, 23] and PPDBv2 [31, 32] using the rigid-body coarse-grained docking tool MAXDo [42]. Both P-262 (262 proteins) and PPDBv2 (168 proteins) cover a large variety of functional classes, such as antibody-antigen, enzyme-regulator, and substrates-inhibitor [11]. For P-262, we efficiently screened 27 millions docking conformations with INTBuilder [10] and the rigidRMSD library [35], and systematically evaluated their quality with respect to the experimentally resolved complex structures available in the Protein Data Bank [3]. For PPDBv2, we obtained the list of acceptable and incorrect conformations from [34]. Among all candidate conformations, we selected 3 902 acceptable or higher quality conformations (L-RMSD < 10.0 Å and I-RMSD < 4.0 Å) and 6 038 incorrect conformations coming from 312 protein pairs for training (**Fig. SI 2**). For about half of these pairs, the docking was performed using the unbound forms of the proteins or their close homologs (≥ 70% sequence identity). As test set, we chose 20 protein pairs not seen during training (**Table SI 1**). For both train and test sets, we reconstructed the high resolution docking conformations with INTBuilder from the Euler angles provided by MAXDo.

#### 2.4.2 BM5: docking conformations produced by HADDOCK

The Docking Benchmark version 5 (BM5) [47] comprises 231 non-redundant target complexes from multiple functional classes, including antibody-antigen and enzyme-inhibitor, and with the corresponding unbound protein structures. We considered a total of 449 158 candidate conformations coming from 142 dimer target complexes. They were generated, selected, and labelled by Renaud and co-authors using the protocol reported in [38]. Specifically, for each target complex, 25 300 docking models were generated using the integrative modelling platform HADDOCK [12] in three stages: (1) rigid-body docking, (2) semi-flexible refinement by simulated annealing in torsion angle space, and (3) final refinement by short molecular dynamics in water [38]. Then, the resulting set of conformations was reduced to avoid redundancy. The conformations with I-RMSD ≤4.0 Å were labelled as near-native. On average, each target complex has ≈230 near-native conformations and ≈2 932 incorrect ones.

#### 2.4.3 Dockground: docking conformations produced by Gramm-X

We downloaded the Dockground database 1.0 [30, 22] from http://dockground.compbio.ku.edu/downloads/unbound/decoy/decoys1.0.zip. This database has 61 target complexes with on average 108 candidate conformations per target generated by the Fast Fourier Transform-based method GRAMM-X [45]. For comparison purpose we followed the experimental setups of GNN-DOVE [48]: in summary, 59 target complexes were chosen and divided into 4 non-redundant groups with respect to the sequence identity (less than 30%) and TM-score [52] (less than 0.5). On average each of these complexes has 9.83 acceptable conformations (L-RMSD ≤5.0 Å) and 98.5 incorrect ones.

### 2.5 Training protocol

We used CCD4PPI to optimize DLA-Ranker hyperparameters. In total, we explored about 10 different architectures by varying the number of convolutional layers, the number of neurons in the fully connected layers, and the dropout rates. We chose the best performing architecture and used it for producing our final results and performing the comparisons with the other methods. We trained several independent models of DLA-Ranker using each of the three considered databases. Using CCD4PPI, we trained 5 models over 20 epochs through a 5-fold cross-validation procedure on the 312 protein pairs (**Fig. SI 3**). For comparison purposes, we reproduced the same training protocols as those reported for DeepRank [38] and GNN-DOVE [48] on BM5 and Dockground, respectively. Specifically, to compare DLA-Ranker with DeepRank, we performed 10-fold cross-validation by splitting the set of 142 dimers selected from BM5 in 114 for training, 14 for validation, and 14 for testing. In total 140 target complexes were used in the test sets (complexes BAAD and 3F1P were not included in the testing). In all three databases the incorrect conformations are much more abundant than the near-native ones. We should stress that, contrary to what was done in [38], we did not augment the input conformational ensemble by random rotations since DLA-Ranker is not sensitive to the orientation of the input conformation. To compare DLA-Ranker with GNN-DOVE, we trained 4 models following 4-fold cross validation. For each model we used 3 non-redundant groups defined from Dockground for training and validation (45 or 44 complexes) and the remaining one for testing (15 or 14 complexes). To compensate the effect of imbalanced training sets and elevate the importance of errors made on near-native poses compared to incorrect ones, we assigned higher weights to the loss of the acceptable class. We used class weights (0.823, 1.273), (0.54, 6.75), and (0.071, 0.929) for CCD4PPI, Dockground, and BM5, respectively.

### 2.6 Evaluation metrics

We used hit rate and enrichment factor to evaluate the performance of DLA-Ranker in ranking candidate conformations. Hit rate curves show the fraction of target complexes in the test set with at least one near-native conformation within the top ranked conformations. Enrichment factor for an individual target complex is defined as the fraction of acceptable conformations found in the top ranked conformations. In case of CCD4PPI we ranked the conformations using a consensus of the 5 trained models. To do so, we first ordered the conformations according to their scores computed from each trained model. Then, we discretised the ranks into 6 bins, namely labels top1, top5, top10, top50, top100, and top200. This way we could represent each conformation as a sequence of ranking labels predicted by 5 models. Finally we “lexicographically” ordered these labels and reported the hit rate of each individual complex separately.

## 3 Results

### 3.1 Identifying near-native conformations

We first assessed DLA-Ranker’s ability to correctly rank candidate conformations. We selected the 1 000 conformations best scored by MAXDo for each of the 20 test protein pairs from CCD4PPI and we re-ranked them according to the *Q* scores predicted by DLA-Ranker. MAXDo evaluates conformations using a physics-based scoring function very similar to that of ATTRACT [51]. For most of the pairs, DLA-Ranker assigned high Q scores to the near-native conformations and discriminated them from the incorrect ones (**Fig. 2A** and **Fig. SI 10**). The top-ranked conformation was near-native in two thirds of the protein pairs (**Fig. 2A**). DLA-Ranker achieved better performance than MAXDo in 11 cases. A particularly difficult case for both MAXDo and DLA-Ranker is the pair 1rkc_A:1ydi_A. Combining DLA-Ranker with the pair potential CIPS [34] allowed enriching the top 200 subset for that pair in near-native conformations, and overall improved the results (**Fig. 2B**). CIPS also improves the performance for the pair 2c9w_A:2jz3_C by surpassing MAXDo and enriching top5. Overall, DLA-Ranker performance do not depend on the extent of conformational change between the docked protein forms and the bound forms (**Fig. 2A-B**, label colors). For instance, one of the cases where it performs very well, the 1ku6_B homodimer, displays a substantial rearrangement (**Fig. 2F**).

**Figure 2:**
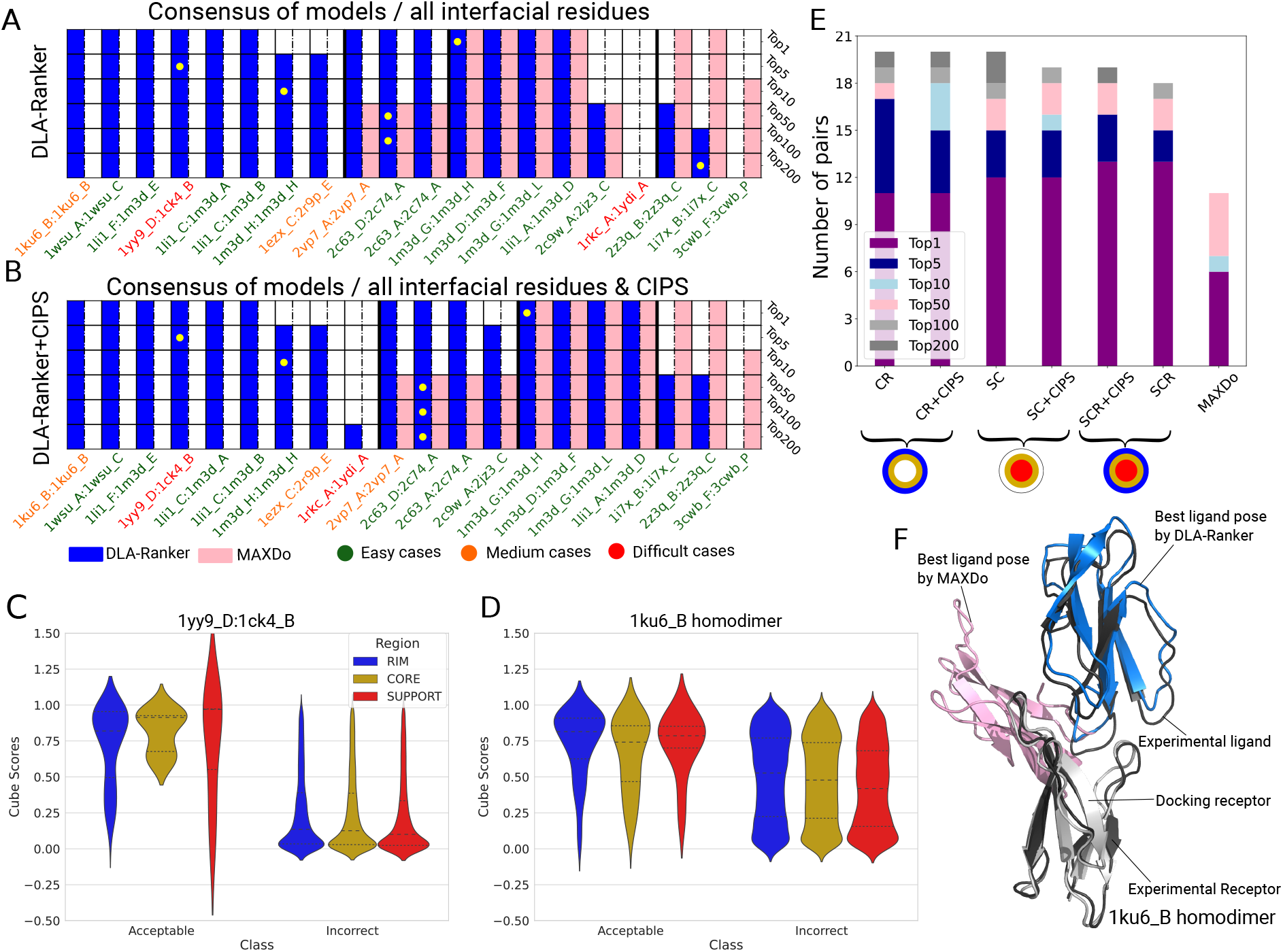
DLA-Ranker performance on CCD4PPI database. **A-B.** Ranking results per protein pair when all interfacial residues are used for train and test according to experimental setup 1 (**Fig. 1C**). For each pair, we report whether some near-native conformations were found in the top 1, 5, …, 200 out of a total of 1 000 conformations generated and selected by MAXDo. A coloured cell indicates the presence of at least one acceptable conformation in the corresponding topX. The pink color corresponds to MAXDo while the blue color corresponds to DLA-Ranker (A) or DLA-Ranker combined with CIPS (B). For each topX, the yellow dot indicates the pair with the highest enrichment factor. The PDB ids are coloured according to the magnitude of the conformational change between the docked forms and the bound forms. Green: none or small. Orange: medium. Red: large. **C-D.** Distribution of individual scores based on S,C,R classes for acceptable and incorrect poses of complex 1yy9_D:1ck4_B (**C**) and 1ku6_B homodimer (**D**). **E.** Comparison between different methods. The SCR, SC, and CR DLA-Ranker models were trained and tested on all interfacial residues, only those in the support and core, or only those in the core and rim, respectively. **F.** Best-ranked candidate conformations for the 1ku6_B homodimer. The reference complex structure is in black, the docked receptor in grey, the ligand conformation selected by MAXDo in pink and that selected by DLA-Ranker in blue.

We further investigated the behaviour of DLA-Ranker for the different sub-regions of the interface, namely the support, core, and rim on two pairs of the database, 1yy9_D:1ck4_B and 1ku6_B homodimer. For both pairs, we observed a wide range of predicted scores within each sub-region (**Fig. 2C-D**). The score distributions for the three subregions often display similar shapes. Nevertheless, it may happen that DLA-Ranker performs significantly differently from one sub-region to the other, as exemplified by the pair 1yy9_D:1ck4_B. In this case, the scores predicted for the residues lying in the support of the interface are not discriminative enough. Averaging the residues’ individual scores over the three interface sub-regions allows correctly classifying the conformations. At the residue level, DLA-Ranker can analyze per-residue scores across near-native conformations to highlight to what extent each residue fits in the interface (**Fig. SI 4A-B**).

### 3.2 Comparison with other scoring functions

We compared DLA-Ranker with two deep learning-based scoring functions, namely DeepRank [38] and GNN-DOVE [48]. We used all interfacial residues for training, and assessed different sub-region combinations (three averaging schemes: SCR, SC, and CR) for testing.

DeepRank applies standard 3D convolutions to a unique voxelised grid representing the interface. On a collection of 10 test sets of 14 target complexes from BM5 (see Methods), DLA-Ranker significantly outperforms DeepRank (**Fig. 3** and **SI 5**). It yields a higher enrichment for both the “raw” conformations produced by the rigid-body docking (**Fig. 3A**) and the semi-flexibly refined conformations (**Fig. 3B**). The enrichment curves obtained on the set of conformations further refined through molecular dynamics simulations in explicit water are almost superimposed (**Fig. SI 5**).

**Figure 3:**
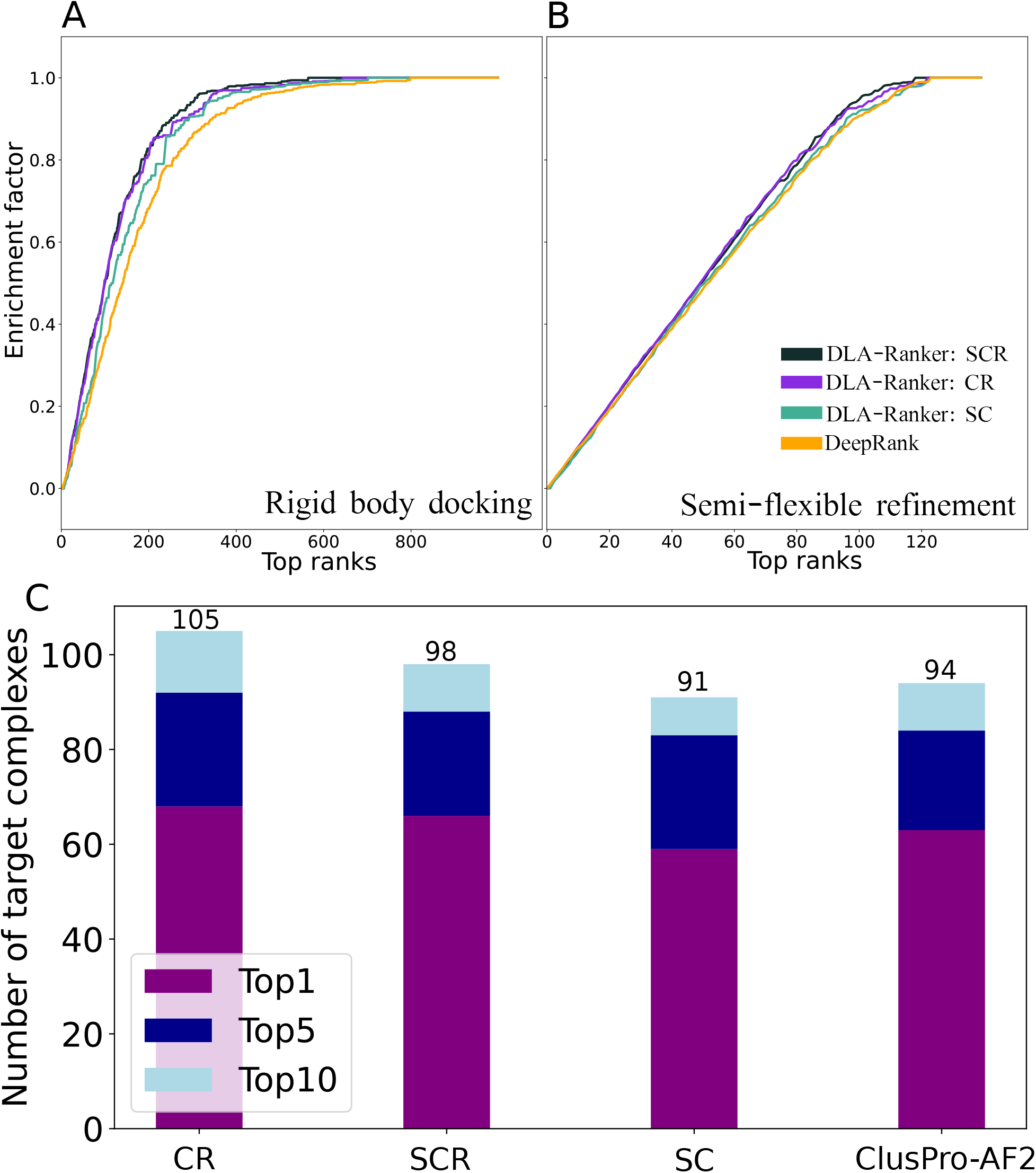
Performance of DLA-Ranker on the 140 dimers of the BM5 database. **A-B.** A comparison between the performance of DLA-Ranker (score averaging schemes SCR, CR, and SC) and DeepRank (orange). Each curve reports the median enrichment over 10 test sets of 14 target complexes (see Methods) See **Fig. SI 5** for both median and the interval between 25% to 75% percentiles. **A.** Only rigid body docking decoys **B.** Decoys with semi-flexible refinement. See **Fig. SI 5** for the performance on decoys with water refinement. **C.** A comparison between combination of HADDOCK and DLA-Ranker and ClusPro-AF2 in protein complex structure prediction in terms of number of target complexes with at least one acceptable or higher quality conformation at top1, top5, and top10.

GNN-DOVE represents the interface as a graph and captures the information on the intermolecular interactions using graph attention mechanisms [48]. DLA-Ranker and GNN-DOVE display comparable hit rates on Dockground (**Fig. SI 6**). While GNN-DOVE identifies a near-native conformation in the top 5 for more complexes than DLA-Ranker, DLA-Ranker covers more complexes when looking at the top 15 conformations. The results differ from one fold to another and this observation may be explained by the small size of the database. It contains about 5 000 conformations versus approximately 10 000 for CCD4PPI and 450 000 for BM5 (see Methods). In the second fold, we observe a lower performance for DLA-Ranker, due to the presence of an outlier complex, namely the ribonuclease inhibitor complex (1DFJ_E_I). The structure of this complex displays several loops on the interface (**Fig. SI 7A**). By comparison, the other structures of the ribonuclease inhibitor complex available in the PDB have more structured interfaces (**Fig. SI 7A**). The t-SNE analysis of the embeddings (averaged over the interface) of 1DFJ_E_I shows less separability compared to those of other complexes from the test set (**Fig. SI 7B-E**).

### 3.3 Influence of the interface description

We investigated whether DLA-Ranker could still discriminate near-native from incorrect conformations with a partial description of the interfaces. To do so, we re-trained DLA-Ranker on CCD4PPI using two different subsets of the interfacial residues: (i) the support and core (SC), or (ii) the core and rim (CR). In the test phase, we aggregated the predicted residue-based scores over the same combination as that used during training (**Fig. 1C**). The results obtained on the 20 test protein pairs from CCD4PPI shows that DLA-Ranker captures sufficient information with a partial description of the interface (**Fig. 2E**). The CR model yielded the best overall performance, and allowed to retrieve near-native conformations in the top 5 for almost all protein pairs (see also **Fig. SI 8**). In addition, we assessed the partial aggregation schemes on BM5 (**Fig. 3** and **SI 5**) and Dockground (**Fig. SI 6**) by the models trained using all interfacial residues. The results are consistent with those on CCD4PPI, with the combination of core and rim yielding a higher performance than the combination of support and core.

We also checked whether we could exploit the topological information of the interface to aggregate the learned residue-based representations. We extracted the embeddings learned by DLA-Ranker on Dockground and used them as node features in a graph representation of the interface (**Fig. 1C**). We observed that the graph-based aggregation does not improve over the global averaging scheme (**Fig. SI 9**C-F). This result can be explained by the fact that the individual embeddings already encode global information about the interface since the labels used during training (acceptable or incorrect) are defined at the level of the interface (**Fig. SI 9A**). This limits the learning capacity of the graph representation, which thus tends to overfit the training set (**Fig. SI 9B**). The similarity between the embeddings in the training set causes homogeneous attention weights and as a result the topology will not influence the learning.

### 3.4 Comparison with ClusPro-AF2

We compared our approach to the recently proposed ClusPro-AF2 protocol [17], where AF2 [20] is used to refine and complement the candidate conformations generated and selected by the docking tool ClusPro [21]. ClusPro-AF2’s overall performance on the test set of 140 dimers from BM5 are similar to those we obtained by applying DLA-Ranker on the candidate conformations produced by HADDOCK (**Fig. 3C**). Moreover, using only the residues located in the core and the rim of the interfaces for DLA-Ranker evaluation increases the number of complexes for which a near-native conformation is found in the top 5 and 10 (**Fig. 3C**, see CR). Considering top 10 ranking, there are 19 complexes for which ClusPro-AF2 predicts acceptable or higher quality conformations while DLA-Ranker cannot find any acceptable one. Five of these complexes (2OT3, 2I9B, 1ATN, 1RKE, 1R8S) have very few acceptable conformations in the ensemble of poses generated by HADDOCK. Reciprocally, there are 23 complexes that are well predicted by DLA-Ranker and are particularly challenging cases for ClusPro-AF2. These include complexes between proteins coming from a pathogen and its host (1EFN, 4H03, 2A9K, 1AK4, 1MAH), complexes from the immune system (1GHQ, 1SBB, 1KXQ, 4M76, 2I25), enzyme-inhibitor complexes (1PXV, 1JTD, 2ABZ), and regulatory complexes (1GLA, 1B6C). While ClusPro-AF2 produces only conformations of very low quality for these complexes, DLA-Ranker is able to identify at least one near-native conformation for 10 of these complexes at top 1, 3 in the top 5, and 2 complexes in the top 10.

### 3.5 Unraveling alternative interfaces

Finally, we explored the potential of DLA-Ranker to discover alternative interfaces. As a case study, we considered the SQD1 enzyme which can self-assemble into homodimers (1qrr) and homotetramers (1i24). We docked the protein (chain 1qrr_A) against itself using ATTRACT and evaluated all interfacial residues detected in the 3 000 best candidate conformations with DLA-Ranker. In **Fig. 4A**, we show the propensity of these residues to have a score higher than 0.5 according to DLA-Ranker. We can clearly identify three patches of residues which appear in acceptable interfaces (**Fig. 4A**, see residues in red). The first one corresponds to the homodimeric interface found in 1qrr (**Fig. 4A**, the other copy of the protein, *i.e*. the partner is in green). The second one corresponds to another interface found in the homotetramer 1i24 (**Fig. 4A**, partner in violet). Finally, the third one is supported by the homotetramer 1wvg, whose chains are homologous to the SQD1 enzyme (E-value=8.58e-4, identified using the PPI3D web server [8]) (**Fig. 4A**, partner in gold). Moreover, the third interface is evolutionary conserved and predicted as an interacting region by JET2Viewer [39] (**Fig. SI 11B**). Altogether, this analysis reveals that DLA-Ranker can be useful to detect multiple binding modes by evaluating individual residues across conformational ensembles. By comparison, looking only at the propensity of each residue to be located at the interface in the candidate conformations [11, 15], without accounting for DLA-Ranker scores, one can clearly identify the first interface but not the two others (**Fig. 4B**). We further compared the ability of ATTRACT and DLA-Ranker in identifying acceptable conformations representative of the different interfaces. ATTRACT and DLA-Ranker (based on the SCR score averaging scheme) find at least one acceptable hit for each of the two first interfaces in the top 22 and 28, respectively. This rank improves to 17 for DLA-Ranker if averaging scheme SC is used (**Fig. SI 11D-E**).

**Figure 4:**
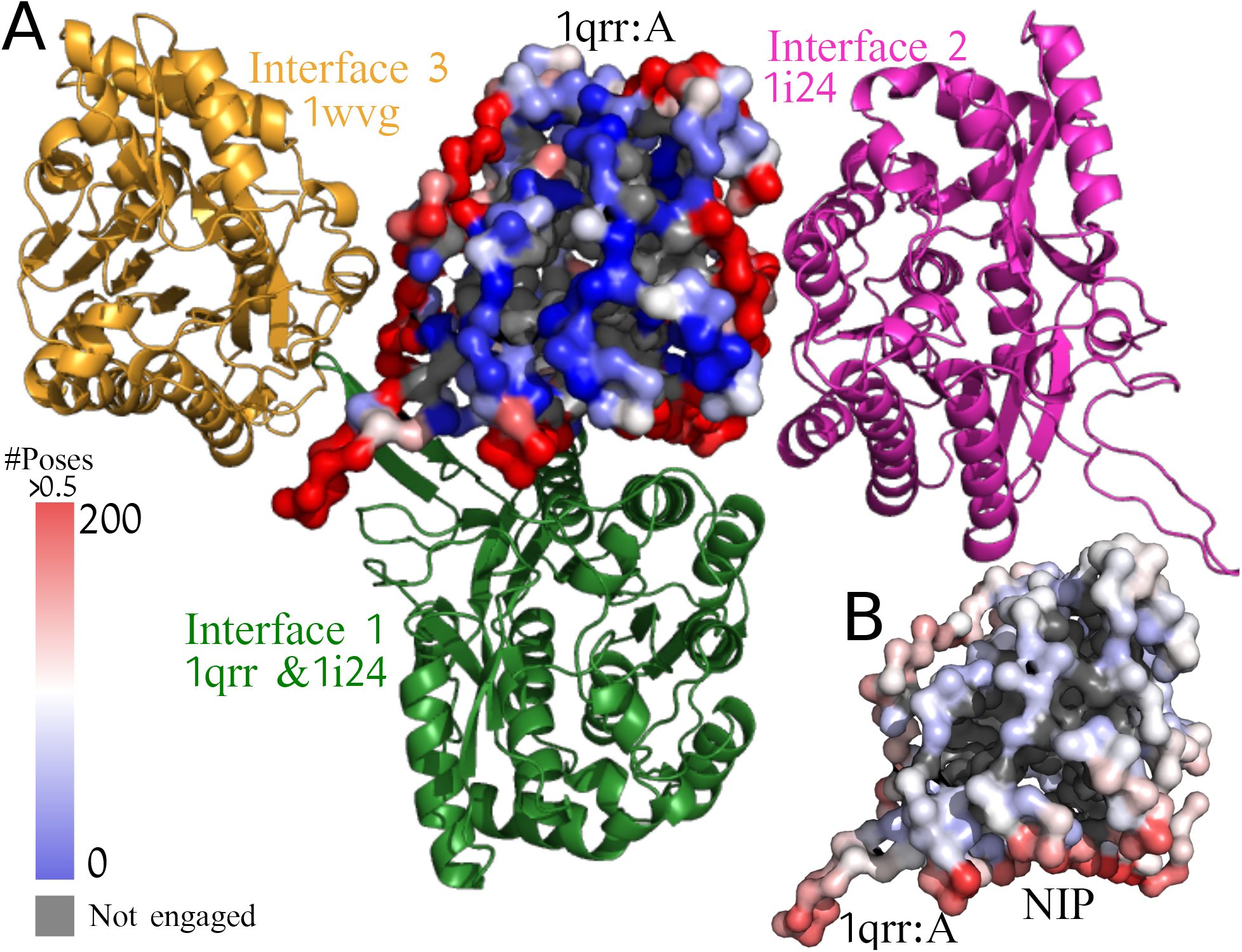
Identification of multiple interaction interfaces for the SQD1 enzyme. **A.** The surface of the protein (chain 1qrr_A) is colored according to the number of conformations (over a total of 3 000) where each residue was found at the interface and was assigned a score higher than 0.5 by DLA-Ranker. Three red patches appear on the surface corresponding to: (i) interface 1 (partner in green, PDB codes: 1qrr, 1i24) (ii) interface 2 (partner in violet, PDB code: 1i24), and (iii) interface 3 (partner in gold, PDB code: 1wvg). **B.** The Normalized Interface Propensity (NIP) shows the tendency of a residue to be part of an interaction site and computed by considering the fraction of docking poses where a residue is found at the interface [11, 15]. It is plotted on 1qrr_A with a color scale going from red (high) to blue (low propensity), and highlights interface 1 but not interfaces 2 and 3, unlike DLA-Ranker.

### 3.6 Runtime and memory usage

The calculations were performed on two GPU clusters: (i) workstations with GPU: NVIDIA GeForce RTX 3090 (24 GB RAM) and CPU: AMD Ryzen 9 5950X and (ii) workstations with GPU: V100 (16 or 32 GB RAM). For a conformation the generation of the interface cubes and their evaluation take on average 1.46 and 0.23 seconds, respectively.

## 4 Discussion

We have shown that it is possible to evaluate complex candidate conformations by learning local 3D atomic arrangements at the interface. We have developed a deep learning-based approach explicitly accounting for the relative orientations of the protein residues while being insensitive to the global orientation of the protein. The method achieves performance better or similar to the state of the art. We obtained the best performance by averaging the per-residue scores predicted over the core and the rim of the interface. DLA-Ranker can be applied to conformational ensembles generated by docking to identify near-native conformations and to discover alternative interfaces. It can be combined with more classical scoring functions. It can also be used to evaluate complexes of any size and is not limited to binary complexes. We envision many applications for the local-environment-based approach of DLA-Ranker, including the identification of physiological interfaces, the discovery of small subsets of cubes dedicated to functional tasks, the construction of phenotypic mutational landscapes, and the prediction of binding affinity.

## Acknowledgements

We thank the Institute for Development and Resources in Intensive Scientific Computing (IDRIS-CNRS) for giving us access to their Jean Zay supercomputer.

## Supplementary Information

**Figure SI 1:**
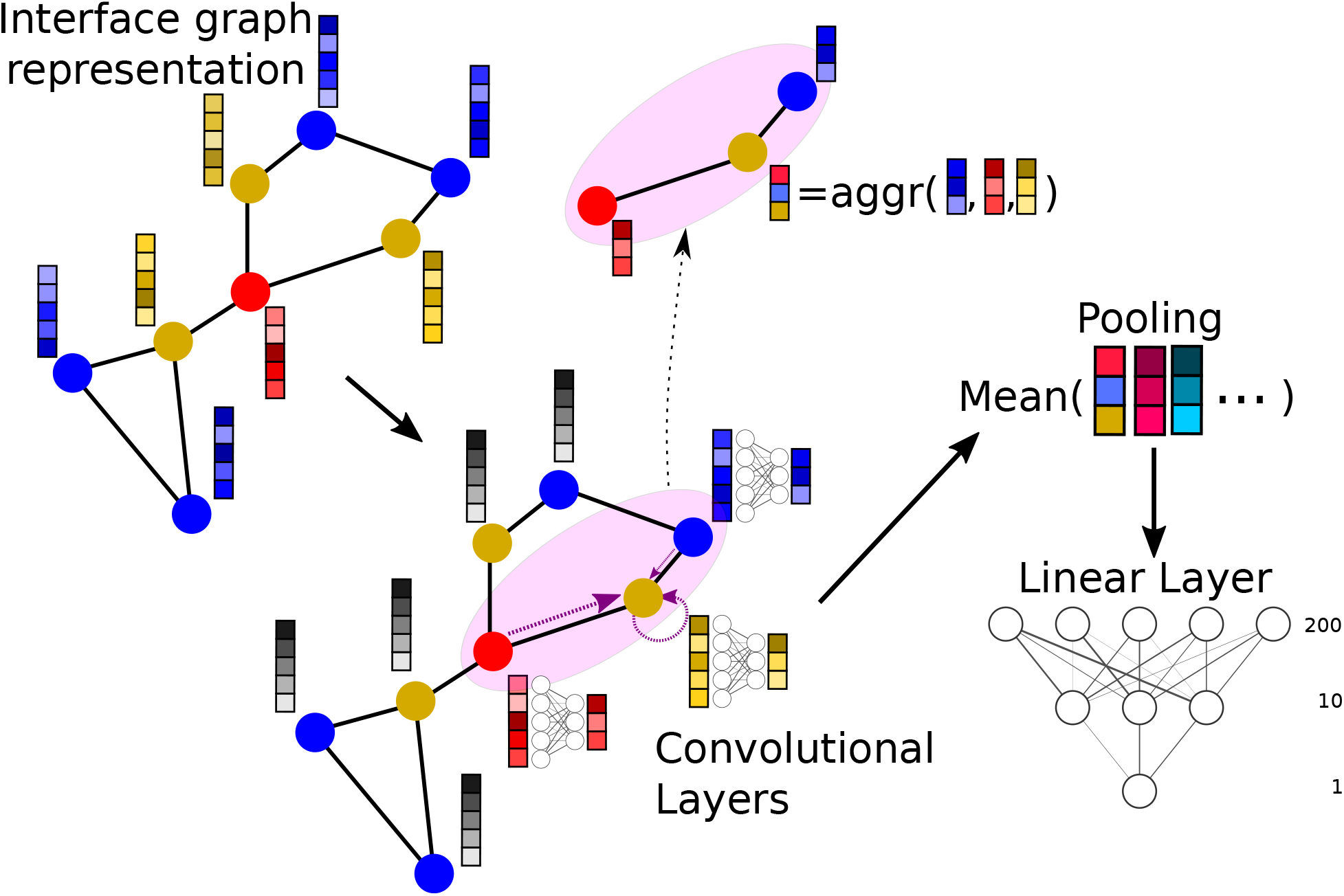
Graph representation of the whole interface. An embedding is extracted for each residue. The structure class of each residue (Support, Core, or Rim) is encoded in this embedding.

**Figure SI 2:**
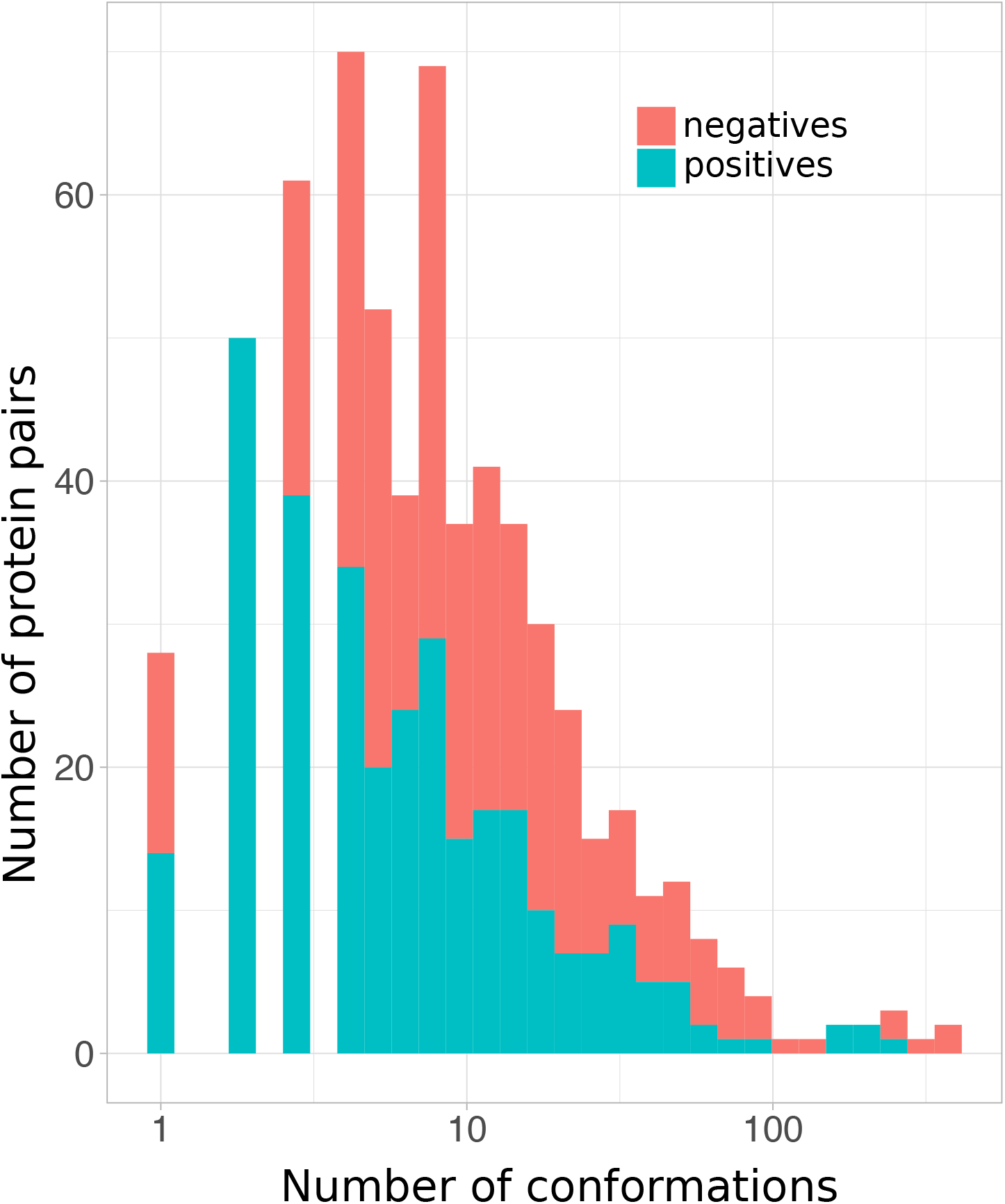
The training set of the CCD4PPI database. Cumulative distributions of positives (acceptable or higher quality conformations) and negatives (incorrect conformations) in the training set of the CCD4PPI database.

**Table SI 1:**
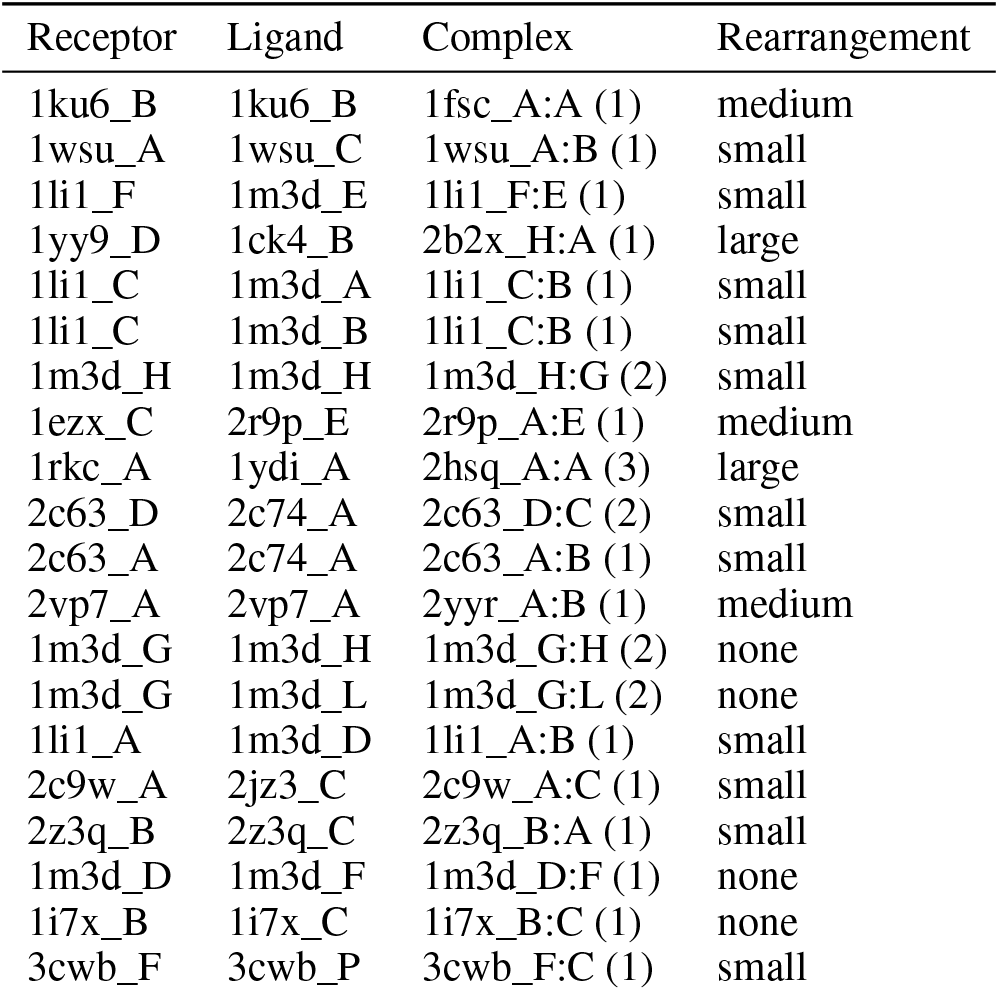
PDB codes for the test set of the CCD4PPI database. The biological assembly ids are given in parenthesis. The extent of conformational rearrangement between the docked protein forms and the bound forms was assessed by the interface root-mean-square deviation (I-RMSD), computed on the C*α* atoms and after superimposition; none: the bound forms were used for docking, small: I-RMSD ≤1.5Å, medium: 1.5Å <I-RMSD < 2.2Å, and large: I-RMSD ≥ 2.2Å.

**Figure SI 3:**
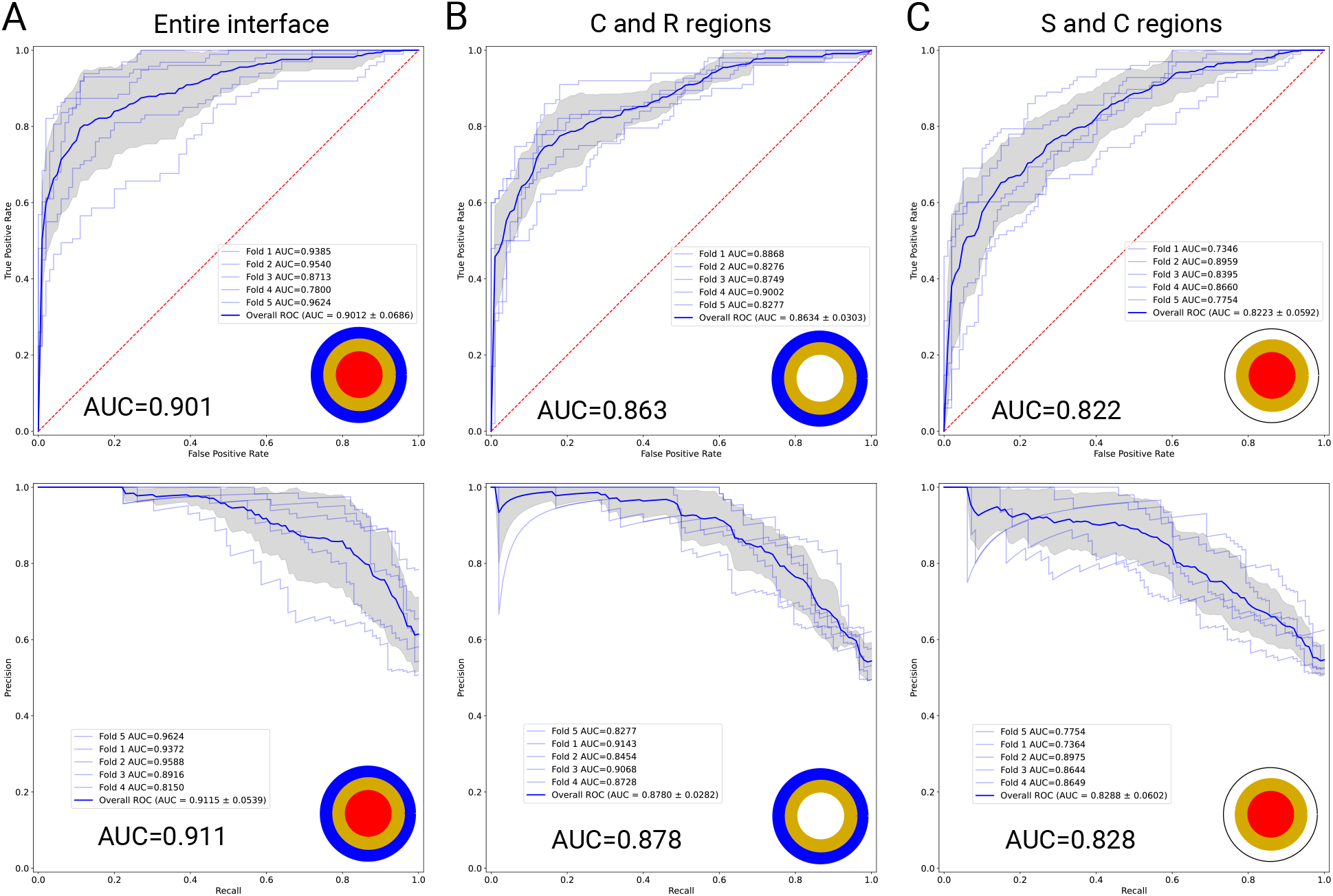
Validation ROC (on top) and PR (at the bottom) curves for 5 × 3 DLA-Ranker models on the CCD4PPI database. The models were trained and validated on entire interfaces (A), on the subset of core and rim interfacial residues (B), or on the subset of support and core interfacial residues (C).

**Figure SI 4:**
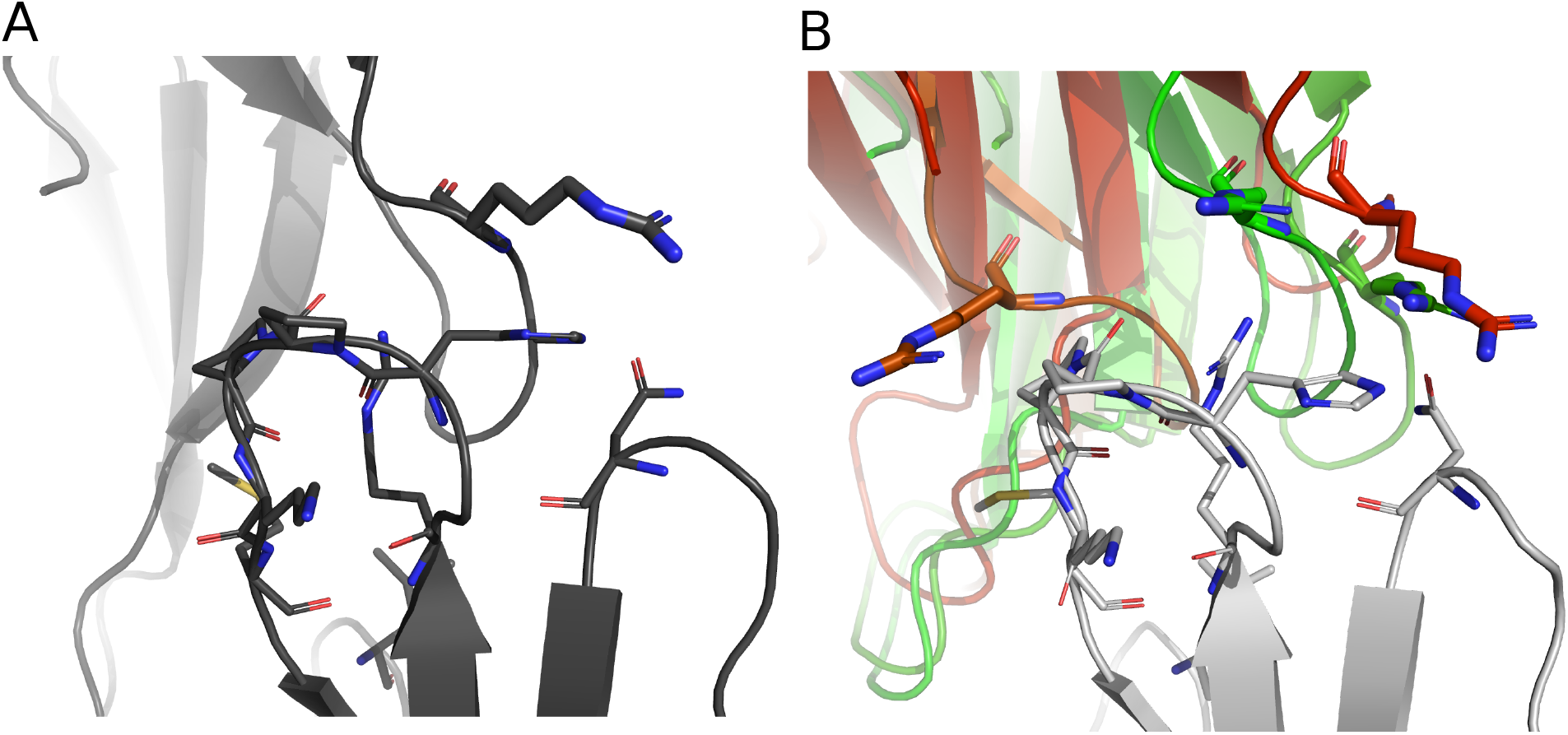
Comparison of individual residue scores between near-native conformations. **A.** Experimental structure of the fasciculin homodimer (PDB code: 1fsc). **B.** Conformations generated by MAXDo using the PDB structure 1ku6_B. The receptor is in gray. The ligand docking poses are colored according to the score predicted by DLA-Ranker for the residue Arg 11, highlighted in sticks. The two poses in green are of medium quality while the red and orange ones are acceptable.

**Figure SI 5:**
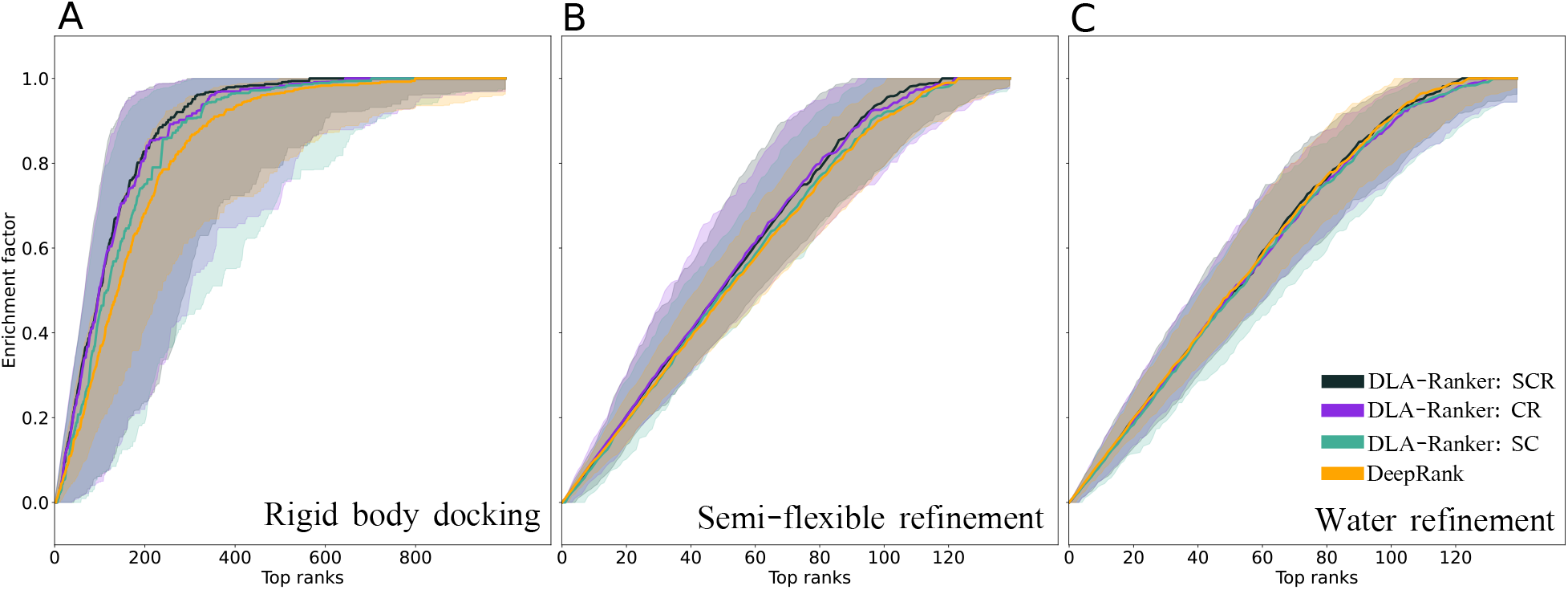
A comparison between the performance of DLA-Ranker (blue) and DeepRank (orange) on the 140 dimers of the BM5 database. The enrichment curves are calculated over 140 target complexes following 10 fold-cross validation. In each fold a model is trained, validated, and tested on 114, 14, and 14 target complexes. The solid line is the median of all individual enrichment curves and the shaded area shows the interval between 25% to 75% percentiles. **A.** Only rigid body docking decoys **B.** Decoys with semi-flexible refinement. **C.** Decoys with water refinement.

**Figure SI 6:**
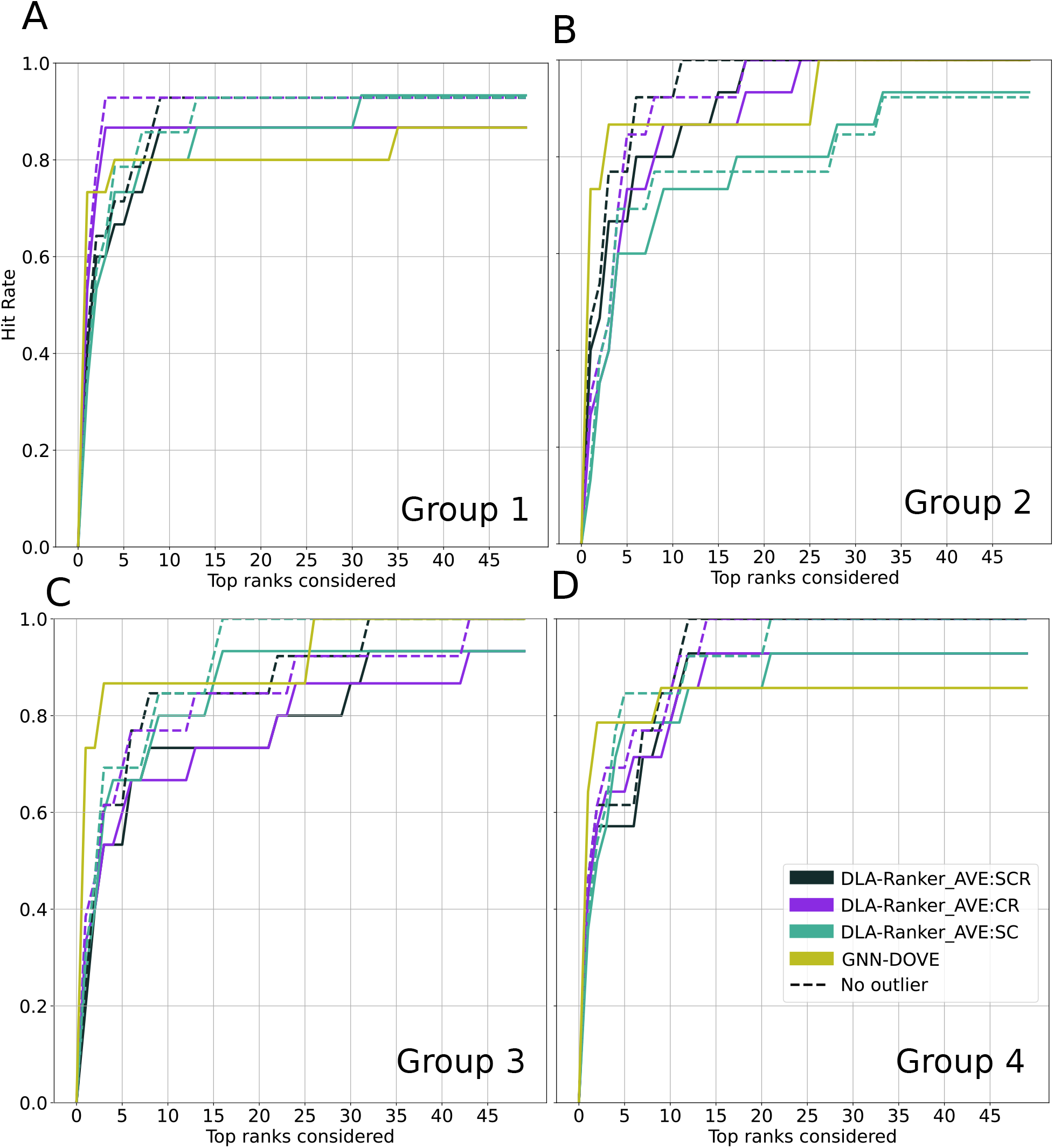
A comparison between the performance of DLA-Ranker and GNN-DOVE on the Dockground database using hit rate curves. **A.** 1 fold 1, a model is trained on groups 2,3,4 and tested on group 1, **B.** fold 2, **C.** fold 3, and **D.** fold 4.

**Figure SI 7:**
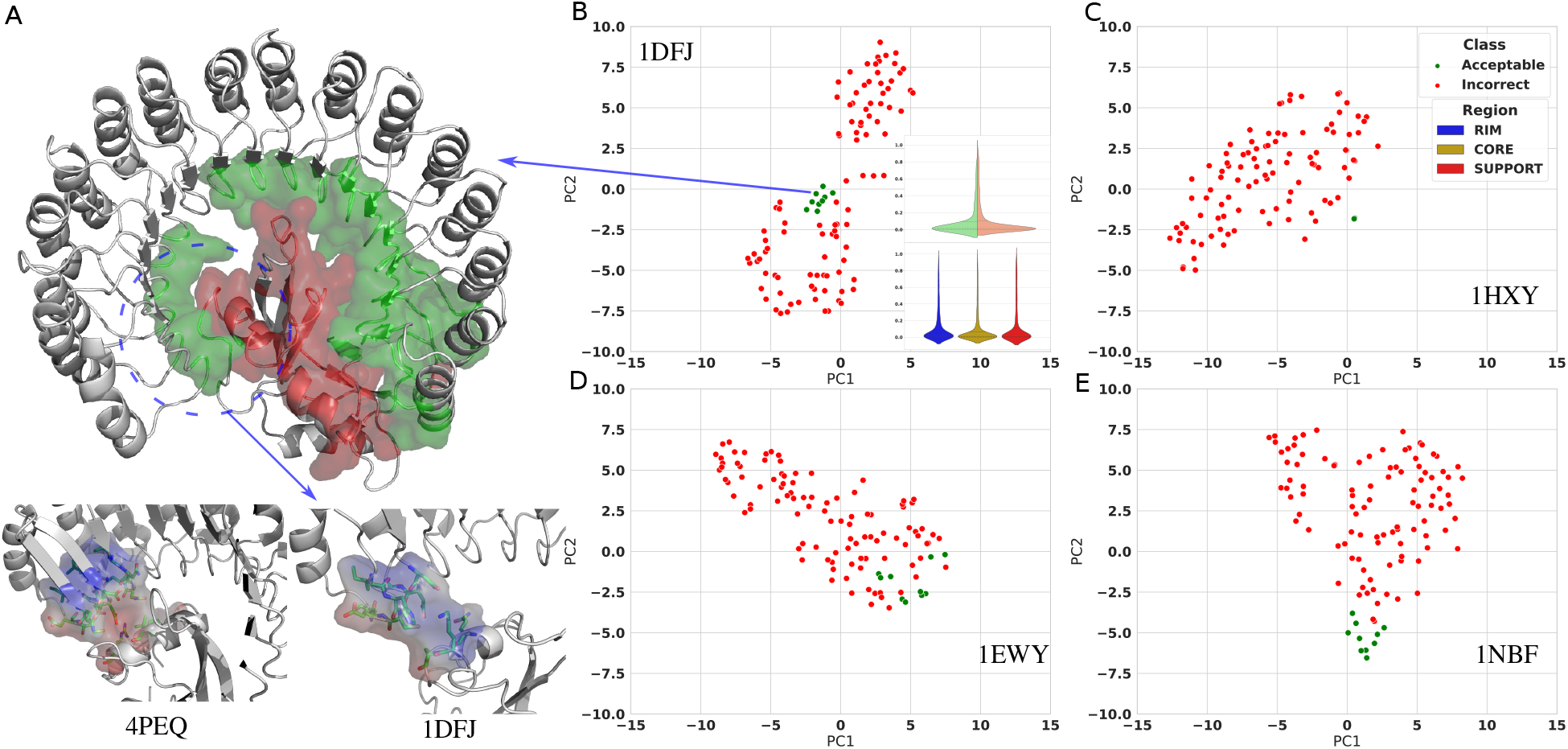
Complex 1DFJ (ribonuclease inhibitor) as an outlier case in the evaluation of a model trained on the Dockground database A. The structure of 1DFJ for Bos taurus with highlighted interaction site. The interaction interface of the inhibitor (green) is mainly composed of loops. The other experimentally resolved structures of ribonuclease inhibitor complexes such as 3TSR (Mus musculus), 4PEQ (Bos taurus), and 4PER (Gallus gallus) have more structured interaction site. Furthermore, the distribution of charges and the physico-chemical properties of the interaction are more favorable (GLN and ASP from inhibitor binds well with ARG from the ribonuclease) for these structures. **B-E** The t-SNE of the embeddings (averaged over the interface) generated by the model on the test set of group 2 for **B.** 1DFJ (inset is the distribution of interfacial residue scores), **C.** 1HXY, **D.** 1EWY, and **E.** 1NBF. Compared to 1DFJ the embeddings of the other complexes are better separated.

**Figure SI 8:**
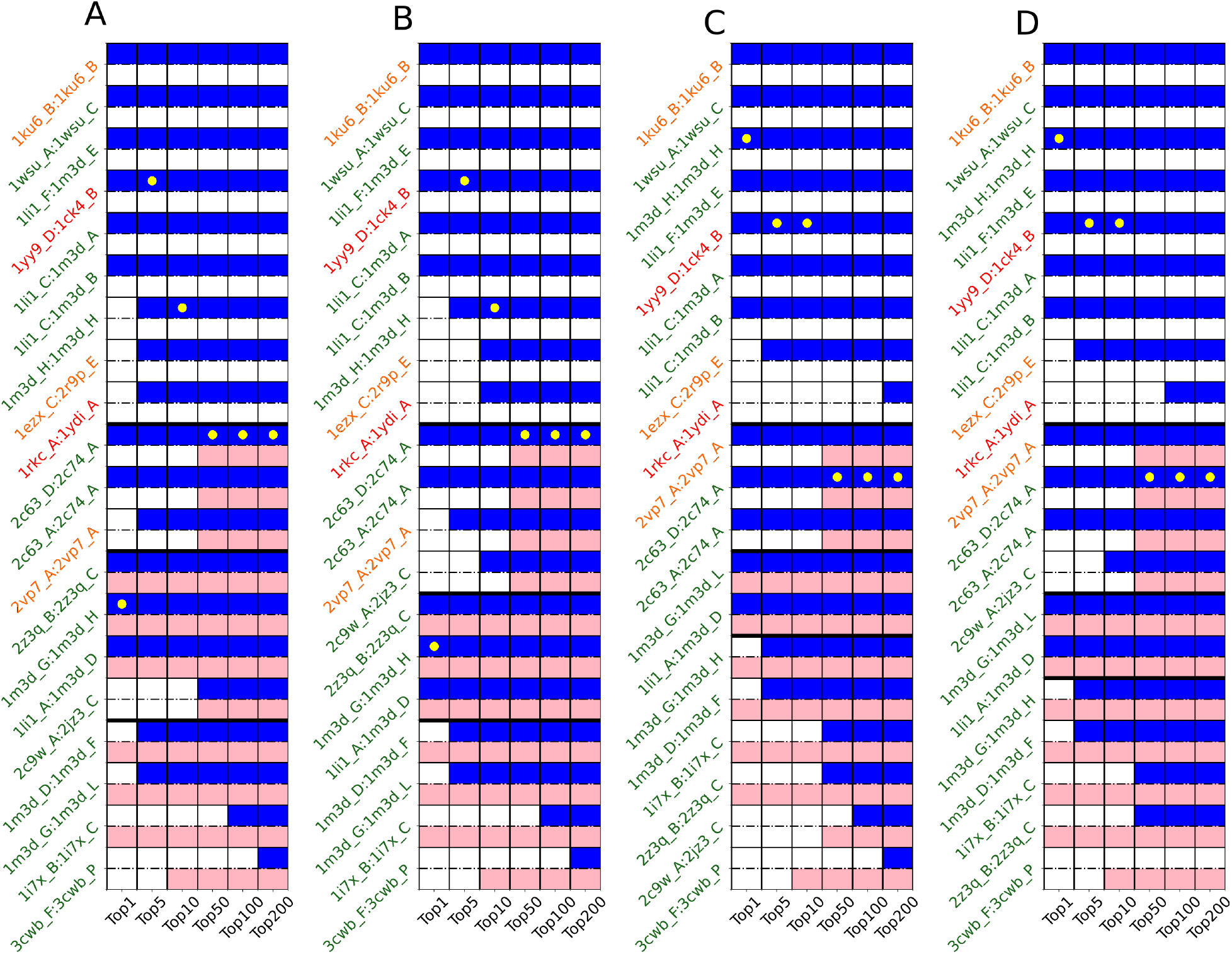
DLA-Ranker performance on CCD4PPI database for different interface descriptions. **A.** DLA-Ranker trained and validated on core and rim residues only (DLA-Ranker-CR). **B.** DLA-Ranker-CR combined with CIPS. **C.** DLA-Ranker trained and validated on support and core residues only (DLA-Ranker-SC). **D.** DLA-Ranker-SC combined with CIPS. The color codes are the same as in Fig. 2.

**Figure SI 9:**
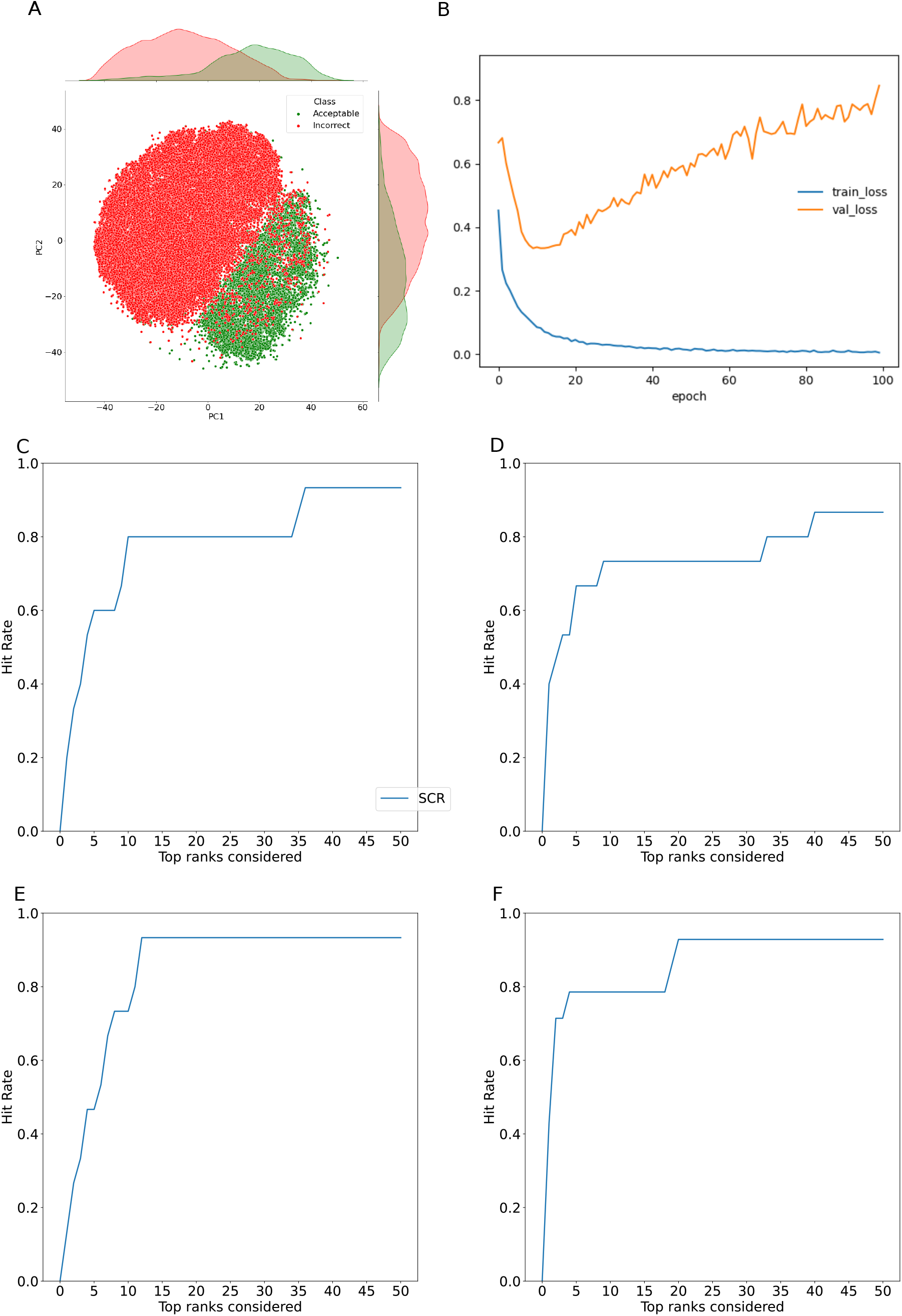
The performance of DLA-Ranker with graph-based aggregation on the Dockground database. **A.** The distribution of the embeddings generated by the model trained on group 3 from Dockground dataset after applying t-SNE. **B.** The training summary of the graph learning on the embeddings. **C-F.** The hit rate curves for 4 test groups of the Dockground database.

**Figure SI 10:**
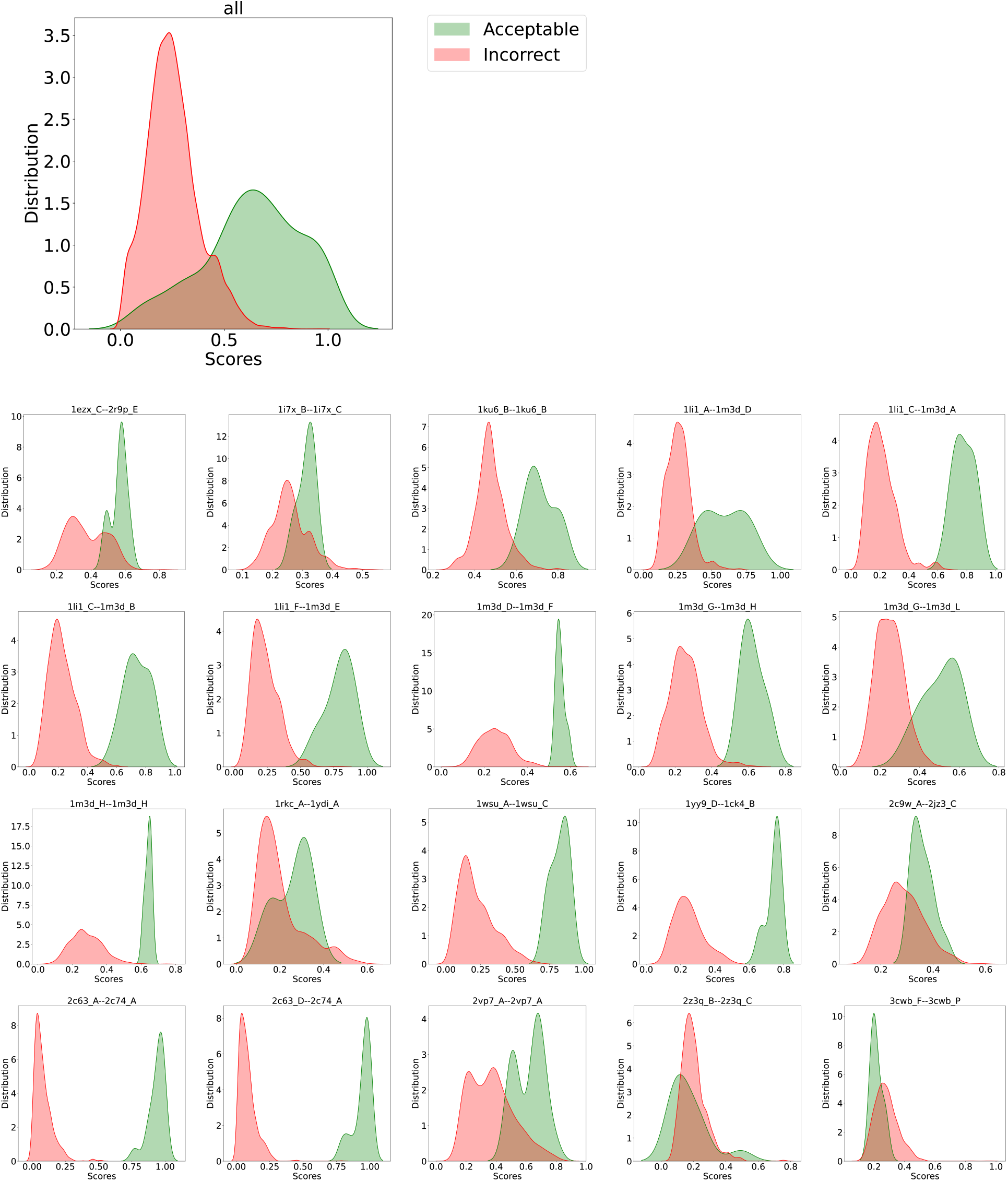
Distributions of DLA-Ranker *Q* scores computed for the test set conformations of the CCD4PPI database. We report the scores computed by one of the 5 DLA-Ranker trained models. The overall distributions are shown on top, and the per protein pair distributions below.

**Figure SI 11:**
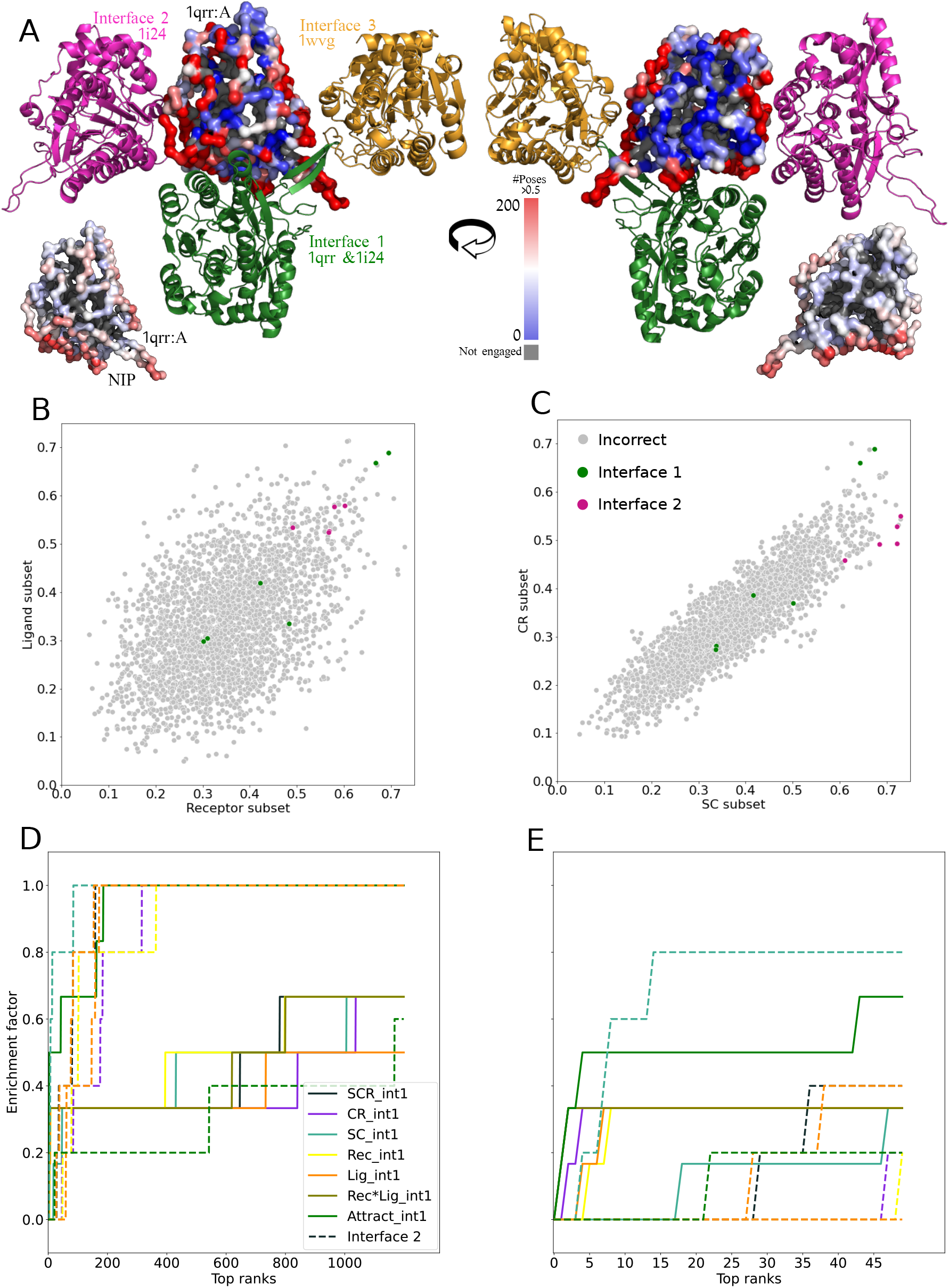
Identification of multiple interaction interfaces for the SQD1 enzyme. **A.** The surface of the protein (chain 1qrr_A) is colored according to the number of conformations (over a total of 3 000) where each residue was found at the interface and was assigned a score higher than 0.5 by DLA-Ranker. Three red patches appear on the surface corresponding to: (i) interface 1 (partner in green, PDB codes: 1qrr, 1i24) (ii) interface 2 (partner in violet, PDB code: 1i24), and (iii) interface 3 (partner in gold, PDB code: 1wvg). The inset is the Normalized Interface Propensity (NIP) showing the tendency of a residue to be part of an interaction site and computed by considering the fraction of docking poses where a residue is found at the interface [11, 15]. It is plotted on 1qrr_A with a color scale going from red (high) to blue (low propensity), and highlights interface 1 but not interfaces 2 and 3, unlike DLA-Ranker. **B-C.** The correlation between scores calculate by selecting residues exclusively from receptor or from ligand (**B**), or residues from SC or CR subests (C). **D-E.** The enrichment curves for DLA-Ranker with different score averaging schemes and for ATTRACT until rank 1200 (**D**) and rank 50 (**E**).

**Figure SI 12:**
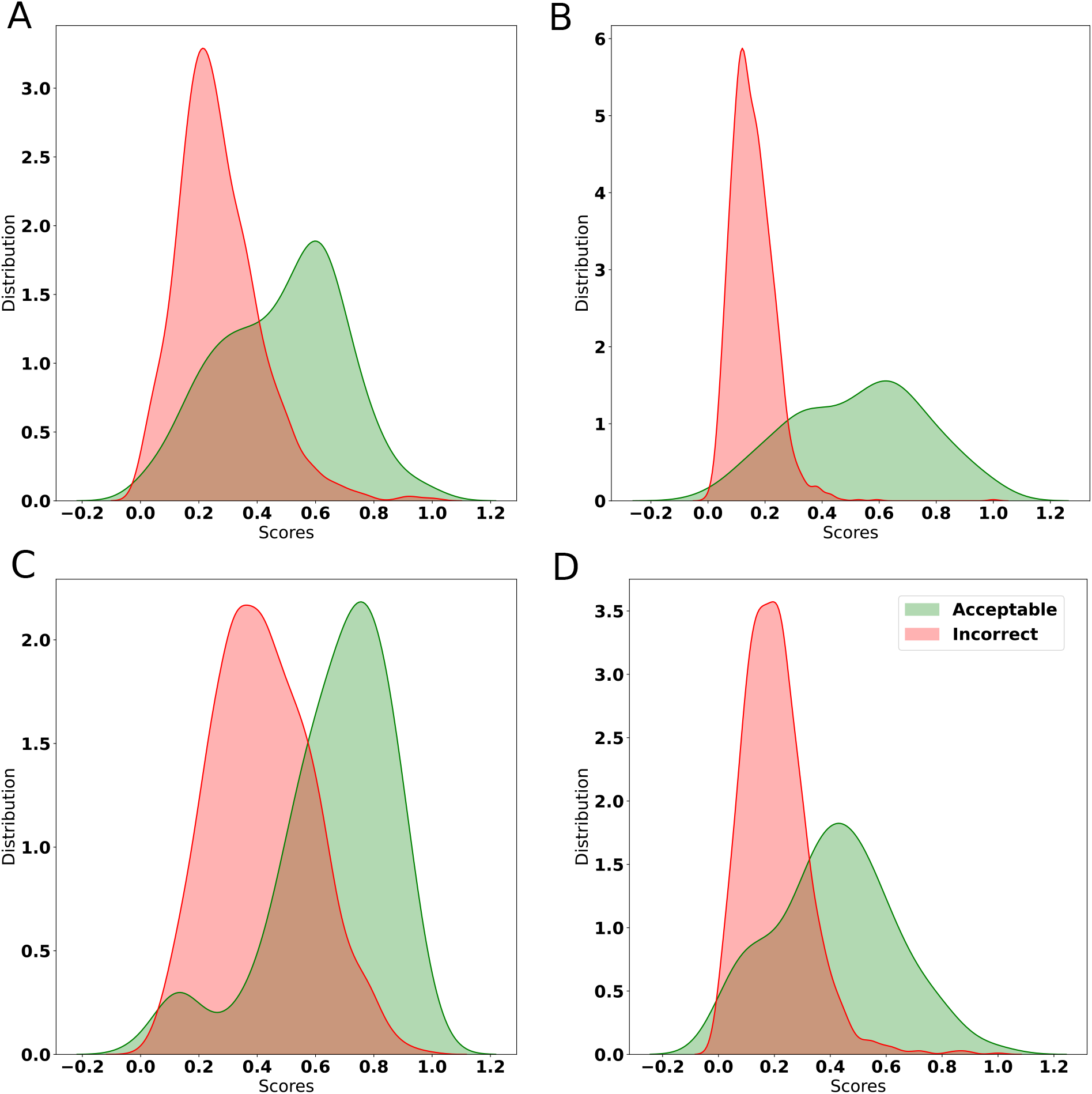
The distribution of scores of 4 test sets from the Dockground database. **A.** group 1, **B.** group 2, **C.** group 3, and **D.** group 4

